# The Function of LncRNA *DRIR* in Freezing Tolerance by Promoting Autophagic Degradation of CP29A and CP29B to Alter Alternative Splicing Patterns of Pre-mRNAs

**DOI:** 10.64898/2026.05.26.727766

**Authors:** Liaoliao Ye, Xiuhua Tang, Jinwei Yang, Zhiquan Qiang, Cun Wang, Liming Xiong, Tao Qin

## Abstract

- Research on the functions and molecular mechanisms of long non-coding RNAs (lncRNAs) involved in regulating plant freezing tolerance is still in its infancy. Our previous research work identified that lncRNA *DROUGHT INDUCED LNCRNA* (*DRIR*) regulates gene expressions in *Arabidopsis*. However, the underlying molecular mechanism is still unknown.
- This study demonstrates that lncRNA *DRIR* regulates plant freezing tolerance by affecting alternative splicing patterns of pre-mRNAs.
- Through chromatin isolation by RNA purification followed by mass spectrometry (ChIRP-MS), we identified two *DRIR* interacting proteins: CP29A and CP29B. We showed that the *drir^D^* mutant, which exhibits elevated *DRIR* expression and *DRIR* overexpression lines showed increased sensitivity to freezing stress, whereas *DRIR* RNAi lines were more tolerant to the stress. CP29A and CP29B bind to nuclear transcripts and, together with *DRIR*, regulate pre-mRNA alternative splicing under freezing stress. Notably, *DRIR* induces the relocalization of CP29A and CP29B to autophagosomes, leading to autophagy-mediated protein degradation.
- Collectively, our findings elucidate the molecular mechanism by which *DRIR* influences the autophagy-based degradation of its binding proteins CP29A and CP29B, thereby regulating plant freezing tolerance by altering the alternative splicing patterns of pre-mRNAs, providing novel insights into the functions and mechanisms of lncRNAs in plants adapting to freezing environments.

## Introduction

Long noncoding RNAs (lncRNAs) are transcripts longer than 200 nt and usually have no obvious protein-coding ability (Gonzales *et al*., 2024). In eukaryotic organisms, lncRNAs have emerged as important regulators participating in nearly every aspect of biology through complex mechanisms, including epigenetic, transcriptional, post-transcriptional, and translational regulation processes (Yu *et al*., 2019; Nojima & Proudfoot, 2022; Kornienko *et al*., 2024). However, compared to lncRNA research in humans and other animal species, the study of plant lncRNAs is still at an early stage, and functional characterization has lagged far behind identification (Palos *et al*., 2023; Kornienko *et al*., 2024).

Cold stress, which includes chilling stress (temperature ranging from 0°C to 15°C) and freezing stress (temperature below 0°C), is one of the major abiotic stresses limiting plant distribution and growth around the world (Huang *et al*., 2023). Protein-coding genes involved in cold stress regulation have been extensively studied, but the roles of lncRNAs in the regulation of cold responses remain largely unknown. Currently, a considerable number of lncRNAs responsive to low temperature have been identified in different plant species, such as *Arabidopsis* (Zhao *et al*., 2018), rice (Yuan *et al*., 2018; He *et al*., 2023), grapes (Wang *et al*., 2019), cassava (Li *et al*., 2021), and *Medicago truncatula* (Zhao *et al*., 2020). One special adaptation process of plants to cold climates is vernalization (Kim *et al*., 2009). During vernalization, the key gene *FLOWERING LOCUS C* (*FLC*), which regulates flowering time, is epigenetically regulated by three lncRNAs: *Cold-induced Long Antisense Intragenic RNA* (*COOLAIR*), *COLD-ASSISTED INTRONIC NON-CODING RNA* (*COLDAIR*) and *Cold of Winter-induced non-coding RNA from the Promoter* (*COLDWRAP*) (Swiezewski *et al*., 2009; Heo & Sung, 2011; Csorba *et al*., 2014; Kim & Sung, 2017). The lncRNA *MADS AFFECTING FLOWERING4 Antisense RNA* (*MAS*) is also involved in vernalization and can activate *MAF4* transcription through epigenetic mechanisms, thereby inhibiting flowering (Zhao *et al*., 2018). In response to cold, the lncRNA *AUXIN-REGULATED PROMOTER LOOP* (*APOLO*) interacts with the transcription factor WRKY42 to activate the expression of *ROOT HAIR DEFECTIVE 6* (*RHD6*) and regulate the growth and development of root hairs under cold stress (Moison *et al*., 2021). Under cold conditions, an lncRNA (MSTRG.13420) is involved in the R-loop formation and regulates the expression of *OsDof16* in rice (He *et al*., 2023). In cassava, the *COLD-RESPONSIVE INTERGENIC LNCRNA 1* (*CRIR1*) enhances the plant’s tolerance to cold stress by increasing the translational yield of stress-responsive genes via COLD SHOCK PROTEIN 5 (MeCSP5) or decreasing DNA methylation (Li *et al*., 2021; Li *et al*., 2024).

Accumulating evidence suggests that lncRNAs play vital roles in plant response to freezing stress. In *Arabidopsis* cold response, the transcription of the *SVALKA*-*asCBF1* lncRNA cascade induces collisions of RNA polymerase II, thereby limiting the expression of *C-repeat/dehydration-responsive element Binding Factor 1* (*CBF1*) and affecting the plant’s freezing tolerance (Kindgren *et al*., 2018). In addition, *SVALKA* recruits PRC2 to the *CBF3* gene locus to ensure the precise progress of cold acclimation (Gómez-Martínez *et al*., 2024). The antisense transcripts of two transcription factors C2H2-TYPE ZINC FINGER FAMILY PROTEIN5 (ZAT5) and B-BOX DOMAIN PROTEIN28 (BBX28) (*asZAT5* and *asBBX28*) in *Arabidopsis* have positive roles in freezing acclimation by priming the sense transcriptions (Meena *et al*., 2024). *COLD INDUCED lncRNA 1* (*CIL1*) is another positive regulator of the plant response to freezing stress in *Arabidopsis* by regulating the expression of multiple stress-related genes during the seedling stage (Liu *et al*., 2022). *Medicago truncatula CBFs intergenic lncRNA2* (*MtCIR2*) positively regulates freezing tolerance by modulating *MtCBF/DREB1s* expression and glycometabolism (Zhao *et al*., 2023). In winter wheat, the lncRNAs *LncR9A*, *lncR117*, and *lncR616* indirectly regulate the *Cu-Zn superoxide dismutase 1* (*CSD1*) expression by competitively binding miR398, improving freezing tolerance (Lu *et al*., 2020). A natural antisense lncRNA transcribed from the antisense strand of *Teosinte branched1/Cycloidea/Proliferating 1* (*DgTCP1*) of chrysanthemum, named *DglncTCP1*, improves chrysanthemum freezing tolerance by activating *DgTCP1* transcription (Li *et al*., 2022). These studies indicate that lncRNAs are important factors in regulating plant responses to freezing environments. However, despite the specific functions of a few lncRNAs regulating plant adaptation to freezing stress being revealed, the detailed molecular mechanisms by which lncRNAs regulate plant freezing stress remain poorly understood.

The *Arabidopsis* genes *CP29A* and *CP29B* are nuclear genes encoding two isoforms of the chloroplast ribonucleoprotein (cpRNP) family (Wang *et al*., 2006). Typically, cpRNPs are abundant in the chloroplast and bind to nascent RNA to form cpRNP-RNA complexes. These cpRNP-RNA complexes confer RNA stability and can regulate RNA maturation, precursor-messenger RNA (pre-mRNA) splicing, or RNA editing (Nakamura *et al*., 2001; Teubner *et al*., 2017; Lenzen *et al*., 2020). There is limited research on the function of CP29B, with existing studies indicating that CP29B is involved in the response to abscisic acid in *Arabidopsis* seeds and suspension cells (Ghelis *et al*., 2008). Research on CP29A has found that CP29A is required for efficient chloroplast RNA processing in response to chilling stress. Under low temperatures, splicing efficiency and stability of chloroplast mRNAs are significantly reduced in the *cp29a* mutants (Kupsch *et al*., 2012; Legen *et al*., 2024; Lenzen *et al*., 2025). A study found that CP29A can also localize to the nucleus and bind to mRNA containing GAN repeat motifs within the nucleus (Gosai *et al*., 2015). The GAN repeat motif has been shown to be related to the regulation of alternative splicing (Wu *et al*., 2014). However, it is still unclear whether CP29A is involved in the regulation of RNA processing within the nucleus.

In our previous study, we identified a lncRNA, *DROUGHT INDUCED LNCRNA* (*DRIR*), which is positively involved in drought and salt stress tolerance in *Arabidopsis* (Qin *et al*., 2017). However, its mechanism of action is still unknown. In this study, we discovered that *DRIR* regulates alternative splicing by affecting autophagy-mediated degradation of CP29A and CP29B, thereby influencing the freezing tolerance of *Arabidopsis* seedlings. This provides new insights for a comprehensive understanding of lncRNA-modulated regulatory mechanisms in plant freezing tolerance through autophagy-dependent protein degradation pathways.

## Materials and Methods

### Plant materials and growth conditions

*Arabidopsis thaliana* plants used were in the Col-0 background. The *drir^D^* mutant (SAIL_813_G12), transgenic lines of *P_DRIR_:GUS*, and the two *DRIR* OE lines have been previously described (Qin *et al*., 2017). The *cp29a* (Salk_003066) and *cp29b* (Salk_008984) mutants were obtained from the Arabidopsis Biological Resource Center. Transgenic lines of *DRIR* RNAi, *CP29A* OE, and *CP29B* OE were generated using the floral dip method. A specific fragment of *DRIR* was inserted into the pFGC5941 vector to produce the RNAi plasmid. The cDNA of *CP29A* or *CP29B* was cloned into the pSuper1300 vector to generate constructs for *35S:CP29A-GFP* and *35S:CP29B-GFP*. The primers used for making these two constructs are described in Table S1. Recombinant plasmids were transformed into *Agrobacterium* strain GV3101. Transgenic plants were generated through *Agrobacterium*-mediated transformation by floral dip. The *DRIR* RNAi *cp29a* and *DRIR* RNAi *cp29b* double mutants were generated by crossing. Seeds were germinated on half-strength Murashige and Skoog (½ MS) medium supplemented with 0.8% agar and 1% sucrose. Plants were grown in a growth chamber under conditions set at 22°C with a 16-h light/8-h dark cycle.

### Histochemical GUS staining

Histochemical GUS staining was conducted using the 5-bromo-4-chloro-3-indolyl-β-D-glucuronic acid (X-gluc). Five-day-old *P_DRIR_:GUS* transgenic plants grown under standard conditions were subjected to -3°C freezing treatment for 1 h. Seedlings with and without the treatment were incubated overnight at 37°C in the GUS staining buffer (0.1 M phosphate buffer at pH 7.0, 1 mM EDTA, 0.05 mM potassium ferricyanide, 0.05 mM potassium ferrocyanide, 1% Triton X-100, and 1 mM X-gluc). After destaining with 70% ethanol and 30% acetic acid, seedlings were observed and photographed using a stereomicroscope.

### Freezing tolerance and electrolyte leakage assays

Two-week-old plants grown on ½ MS plates under normal conditions were subjected to -4°C freezing treatment for 1 h. For cold acclimation, plates were transferred to a 4°C chamber for 4 days, and then plants were subjected to -6°C freezing treatment for 0.5 h. The freezing program was set at 0°C and decreased by 1°C per hour to indicated temperatures. After the freezing treatment, the electrolyte leakage was measured. For survival rate assay, the plates were transferred to 4°C for 12 h in darkness after freezing treatment, then moved to normal conditions for a 3-day recovery.

### Chromatin isolation by RNA purification followed by mass spectrometry

Chromatin isolation by RNA purification followed by mass spectrometry (ChIRP-MS) was performed as previously described with modifications (Rigo *et al*., 2020). Seven-day-old *DRIR* OE seedlings were cross-linked with 3% formaldehyde and then homogenized in Honda buffer (0.44 M Sucrose, 1.25% Ficoll, 2.5% Dextran T40, 10 mM MgCl_2_, 20 mM HEPES at pH 7.4, 5 mM DTT, 1% Triton X-100, 1 mM PMSF, 1 × Protease inhibitor cocktail, and 2 U/ml RNase inhibitor). After filtering through two layers of Miracloth, the solution was centrifuged at 3,000 *g* for 15 min at 4°C. The pellets were washed three times with Honda buffer, resuspended in nuclear lysis buffer (50 mM Tris–HCl pH 7.5, 2 mM MgCl_2_, 10 mM EDTA, 1 mM DTT, 1% SDS, 0.1 mM PMSF, 1 × Protease inhibitor cocktail, 0.1 U/µl RNase Inhibitor), and sonicated by sonicator QSONICA Q700. After centrifugation at 12,000 *g* at 4°C for 10 min, the supernatant was diluted with hybridization buffer (50 mM Tris pH 7.5, 750 mM NaCl, 1% SDS, 1 mM EDTA, 15% formamide, and 1 × Protease inhibitor cocktail). The mixture was hybridized with biotinylated probe against *DRIR* (5’-CTCCAAACTCCTTTATTTCTTAACCAAAAGTTACAATTCATGAGAAGATGATCTAGAACATCATTTCTAGACTCATCTTCTAAATCTCACACACGAGATTGTTTACACAAATTGCATAAAGCTCTCTAAACAATGAGAGTACCTATTTATAACCAAAAAGCAGTAAAAGATAGATGCGGATATTACCTCAGAATATCTTC-3’) or biotinylated scramble probe (5’-GTGTAACACGTCTATACGCCCAGTGTAACACGTCTATACGCCCAGTGTAACACGTCTATACGCCCAGTGTAACACGTCTATACGCCCAGTGTAACACGTCTATACGCCCAGTGTAACACGTCTATACGCCCAGTGTAACACGTCTATACGCCCAGTGTAACACGTCTATACGCCCAGTGTAACACGTCTATACGCCCAGT-3’) at 50°C for 4 h. Then 100 µl of Streptavidin Magnetic Beads (Beyotime Biotechnology) was added to each sample and incubated at 50°C for 1 h. Captured beads were washed three times with high-salt wash buffer (2 × SSC, 0.5% SDS, 1 mM DTT, and 1 mM PMSF, 1 × Protease inhibitor cocktail, and 2 U/ml RNase inhibitor). Co-purified proteins were eluted and RNase treated according to the manufacturer’s protocol (Thermo Scientific). Then, 1.8 ml of TCA acetone (5 ml 6.1 N TCA + 45 ml acetone + 35 µl β-mercaptoethanol) was added into the samples and incubated overnight at 80°C. Following washing with acetone wash buffer (120 ml acetone, 84 µl β-mercaptoethanol), pellets were solubilized in trypsin buffer (Promega) for digestion into small peptides and subjected to mass spectrometry analysis by a Q ExactiveTM HF hybrid quadrupole-Orbitrap mass spectrometer (Thermo Scientific). Data were analyzed using Mascot software (Matrix Science).

### RNA-EMSA

The cDNAs of CP29A, CP29B, and GRP7 were cloned into the pGEX4T-1 vector and recombinant proteins of GST, GST-CP29A, GST-CP29B, and GST-GRP7 were purified. Specific primers used are shown in Table S1. *DRIR* RNAs were synthesized in vitro with or without biotin-labelled UTP according to the HiScribe® SP6 RNA Synthesis Kit (NEB). The RNA electrophoretic mobility shift assay (RNA-EMSA) was performed using the Light Shift Chemiluminescent RNA EMSA Kit (Beyotime Biotechnology). Briefly, the labelled RNA and the recombinant proteins were incubated for 30 min at room temperature in a 10 μl reaction system (1 × REMSA Binding Buffer, 2 μg protein, 0.5 nM labelled RNA, 1 U/µl RNase Inhibitor, and unlabeled RNA). The mixtures were subjected to a 6% native polyacrylamide gel electrophoresis in 0.5× TBE buffer, and then RNAs were transferred onto a nylon membrane. After UV crosslinking, the labelled RNA on the membrane was detected using a Chemiluminescence Biotin Labeled Nucleic Acid Detection Kit (Beyotime Biotechnology).

### RNA-protein pull-down assay

Biotin-labeled *DRIR* RNAs were synthesized in vitro and GST, GST-CP29A, GST-CP29B recombinant proteins were purified as described above. One microgram of biotin-labeled RNAs and 2 μg of recombinant proteins were mixed in pull-down buffer (1 × REMSA Binding Buffer, 1% NP-40, 1 U/µl RNase Inhibitor) and incubated at room temperature for 30 min. Fifty microliters of washed Streptavidin Magnetic Beads were added to the reaction system and incubated for another 2 h at 4°C. The beads were washed briefly five times using binding buffer and then boiled in SDS buffer. The supernatant was analyzed by immunoblotting using an anti-GST antibody.

### Isothermal titration calorimetry assay

All isothermal titration calorimetry (ITC) experiments were performed on a Nano ITC CV (Waters Technologies Corporation) at 20°C. *DRIR* RNAs were synthesized in vitro and GST, GST-CP29A, GST-CP29B recombinant proteins were purified. Before titration, all *DRIR* RNAs and recombinant proteins were dialyzed in PBS buffer (137 mM NaCl, 2.7 mM KCl, 10 mM Na_2_HPO_4_, 2 mM KH_2_PO_4_). A final concentration of 150 μM *DRIR* RNAs was titrated into 10 μM GST, GST-CP29A, or GST-CP29B protein by 24 serial injections of 2 µl with 180 s intervals between injections with 300 rpm stirring speed. Using PBS buffer for RNA titration as the blank control, independent and blank fitting were performed using NanoAnalyze software to calculate thermodynamic parameters including stoichiometry (N), dissociation constant (K_D_), enthalpy change (ΔH), and entropy change (ΔS).

### RIP-qPCR assay

RNA immunoprecipitation assay followed by qPCR (RIP-qPCR) assays were performed as previously described with modification (Rigo *et al*., 2020). Five grams of WT, *35S:CP29A-GFP*, or *35S:CP29B-GFP* seedlings were cross-linked with 1% (w/v) formaldehyde for 15 min under vacuum, followed by termination with 125 mM glycine. The nuclei were then isolated and lysed as described above in ChIRP-MS section. The nuclear extract (input) was incubated with 100 µl of anti-GFP magnetic beads (Beyotime Biotechnology) at 4°C for 4 h. Beads were subsequently washed with buffer 1 (20 mM Tris-HCl pH 8.1,150 mM NaCl, 1% Triton X-100, 2 mM EDTA, 20 U/ml RNase inhibitor), buffer 2 (20 mM Tris-HCl pH 8.1, 500 mM NaCl, 1% Triton X-100, 2 mM EDTA, 20 U/ml RNase inhibitor), buffer 3 (0.25 M LiCl, 1% Sodium deoxycholate, 10 mM Tris-HCl pH 8.1, 1% Igepal CA-630, 1 mM pH 8.0 EDTA, 20 U/ml RNase inhibitor), and buffer 4 (10 mM Tris-HCl pH 8.1, 1 mM EDTA, 20 U/ml RNase inhibitor). Following Proteinase K (Beyotime Biotechnology) treatment, RNA was extracted and purified using TRIzol. Ten microliters of nuclear extracts were used for input RNA extraction. Quantitative real-time PCR (qRT-PCR) reactions were performed using specific primers to determine the enrichment of target RNAs. Specific primers used are shown in Table S1.

### The identification and validation of AS events

Six-day-old WT, *drir^D^*, *DRIR* OE, *cp29a*, and *cp29b* seedlings were subjected to -3°C freezing treatment for 1 h and then harvested for total RNA extraction. High-quality RNA samples with RIN > 7.0 were used to construct the sequencing library and subjected to the 2 × 150 bp paired-end sequencing on an Illumina Novaseq™ 6000 (LC-Bio Technology CO., Ltd., Hangzhou, China). Following filtering by Cutadapt (https://cutadapt.readthedocs.io/en/stable/, version: cutadapt-1.9), reads were mapped to the *Arabidopsis* reference genome (https://www.arabidopsis.org/) using Hisat2 (v2-2.2.1). The transcript abundance was estimated using FPKM values (Fragments Per Kilobase of transcript sequence per Million mapped fragments). rMATS v4.1.1 (http://rnaseq-mats.sourceforge.net) was used to identify alternative splicing events and analyze differential alternative splicing (AS) events between WT and other samples. AS events with a false discovery rate (FDR) < 0.05 in a comparison were identified as significant. Differential AS events were identified with the option: FDR < 0.05, IncLevelDifference ≥0.1 or ≤ 0.1.

To validate the differential AS events, total RNAs of stress treated WT, *drir^D^*, *DRIR* OE, *cp29a*, or *cp29b* seedlings were extracted with RNeasy Plant Mini Kit following the manufacturer’s protocols (QIAGEN). DNase-free RNA was reverse-transcribed using HiScript II Q RT SuperMix Kit (Vazyme). qRT-PCR or reverse transcription PCR (RT-PCR) was used to determine the level of alternative spliced or unspliced RNAs. The splicing ratio is defined as the ratio of spliced RNA level to the level of unspliced RNA. Splicing-specific primers are presented in Table S1.

### Microscopy observation

To observe subcellular localization, constructs of *35S:CP29A-GFP* or *35S:CP29B-GFP* were introduced into tobacco leaves with or without *35S:DRIR* via *Agrobacterium*-mediated transformation. Alternatively, *Arabidopsis* protoplasts of WT, *drir^D^*, and *DRIR* OE were transiently transformed with *35S:CP29A-GFP* or *35S:CP29B-GFP* via polyethylene glycol (PEG)-mediated transformation according to standard protocols (Yoo *et al*., 2007). Confocal images were acquired using a Zeiss LSM900 or an Olympus FV3000 confocal microscope. Fluorescence signals were visualized using a 488 nm laser, and emission was collected between 500-525 nm, while the emission wavelength for chloroplast auto-fluorescence was above 610 nm. For the observation of co-localization of CP29A and CP29B with autophagosomes, constructs of *35S:CP29A-GFP* or *35S:CP29B-GFP* were introduced into tobacco leaves together with *35S:DRIR* and *35S:ATG6-RFP*. Infiltrated leaves were thoroughly smeared with either 10 mM 3-Methyladenine (3-MA) or 0.1% DMSO 12 hours prior to imaging. The excitation wavelength for RFP was 532 nm, and emission bandwidth was between 560 and 610 nm.

### Subcellular fractionation and immunoblot assay

Two grams of seedlings of *35S:CP29A-GFP* or *35S:CP29B-GFP* in WT or *DRIR* OE background were ground and homogenized in 4ml of Honda buffer (0.44 M sucrose, 1.25% Ficoll 400, 2.5% Dextran T40, 20 mM HEPES pH 7.4, 10 mM MgCl_2_, 0.5% Triton X-100, 5 mM DTT, 1 mM PMSF, and protease inhibitor cocktail). After filtering through two layers of Miracloth, 300 μl of the solution was removed as total protein fraction, and the remaining filtrates were centrifuged at 3,000 *g* for 15 min at 4°C. Nuclear pellets were washed three times with Honda buffer and the final pellets were resuspended in 300 μl of Honda buffer. The protein content of CP29A-GFP or CP29B-GFP in total proteins or nuclear fractions was assessed using an anti-GFP antibody. Histone H3 was used as an internal control for nuclear fractions and a loading control while Rubisco was used as a loading control for total protein. For MG132, seedlings of *35S:CP29A-GFP* or *35S:CP29B-GFP* in WT or *DRIR* OE background were treated with 0, 20, or 50 µM MG132 for 4 h. For chloroquine treatment, the seedlings were exposed to 200 µM chloroquine for 0, 2, or 4 h, respectively. After treatment, total proteins and nuclear proteins were extracted, and the protein content of CP29A-GFP or CP29B-GFP was assessed.

## Results

### *DRIR* interacts with CP29A and CP29B

Our previous research revealed that the lncRNA *DRIR* is primarily localized in the nucleus. It may influence the expression of stress-responsive genes through a nuclear regulatory mechanism. However, the specific molecular mechanisms remain unclear (Qin *et al*., 2017). In the present study, to elucidate the regulatory mechanism of *DRIR*, we performed chromatin isolation by RNA purification followed by mass spectrometry (ChIRP-MS) to identify the potential *DRIR*-binding proteins. Using the *DRIR*-specific probe from our previous fluorescence in situ hybridization study (Qin *et al*., 2017), we identified five proteins, with MPK4 and SESA2 also appearing in the negative control (scrambled probe) sample. The remaining three proteins were all RNA-binding proteins, among which CP29B showed the highest protein sequence coverage (Fig. 1a). To verify the binding characteristics of these three proteins with *DRIR*, we performed RNA electrophoretic mobility shift assay (RNA-EMSA) to examine their interactions in vitro. The results demonstrated that GST-CP29A and GST-CP29B could specifically bind to *DRIR*, while the GST tag control showed no binding activity (Fig. 1b,c). Intriguingly, no interaction was detected between GLYCINE RICH PROTEIN 7 (GRP7) and *DRIR* (Fig. S1), probably because the interaction is indirect or due to other unknown factors. To further confirm the interaction of CP29A and CP29B with *DRIR*, we conducted RNA-protein pull-down and isothermal titration calorimetry (ITC) assays. In the RNA-protein pull-down experiment, both GST-CP29A and GST-CP29B were pulled down by biotin-labeled *DRIR* RNA, whereas the GST tag alone was not (Fig. 1d). In the ITC assay, significant temperature changes were observed when titrating *DRIR* RNA into GST-CP29A and GST- CP29B samples. Indeed, measurable interactions in the nanomolar range were detected between *DRIR* and CP29A or CP29B (Fig. 1e,f). In order to validate the interaction of these two proteins with *DRIR* in vivo, we performed a RNA immunoprecipitation assay followed by qPCR (RIP-qPCR) on nuclear extracts. The results showed that *DRIR* was specifically enriched in the immunoprecipitates of *35S:CP29A-GFP* and *35S:CP29B-GFP* using anti-GFP magnetic beads, while the negative control *ACTIN2* was not enriched (Fig. 1g). These results indicate that CP29A and CP29B can specifically bind to *DRIR*.

**Fig. 1.**
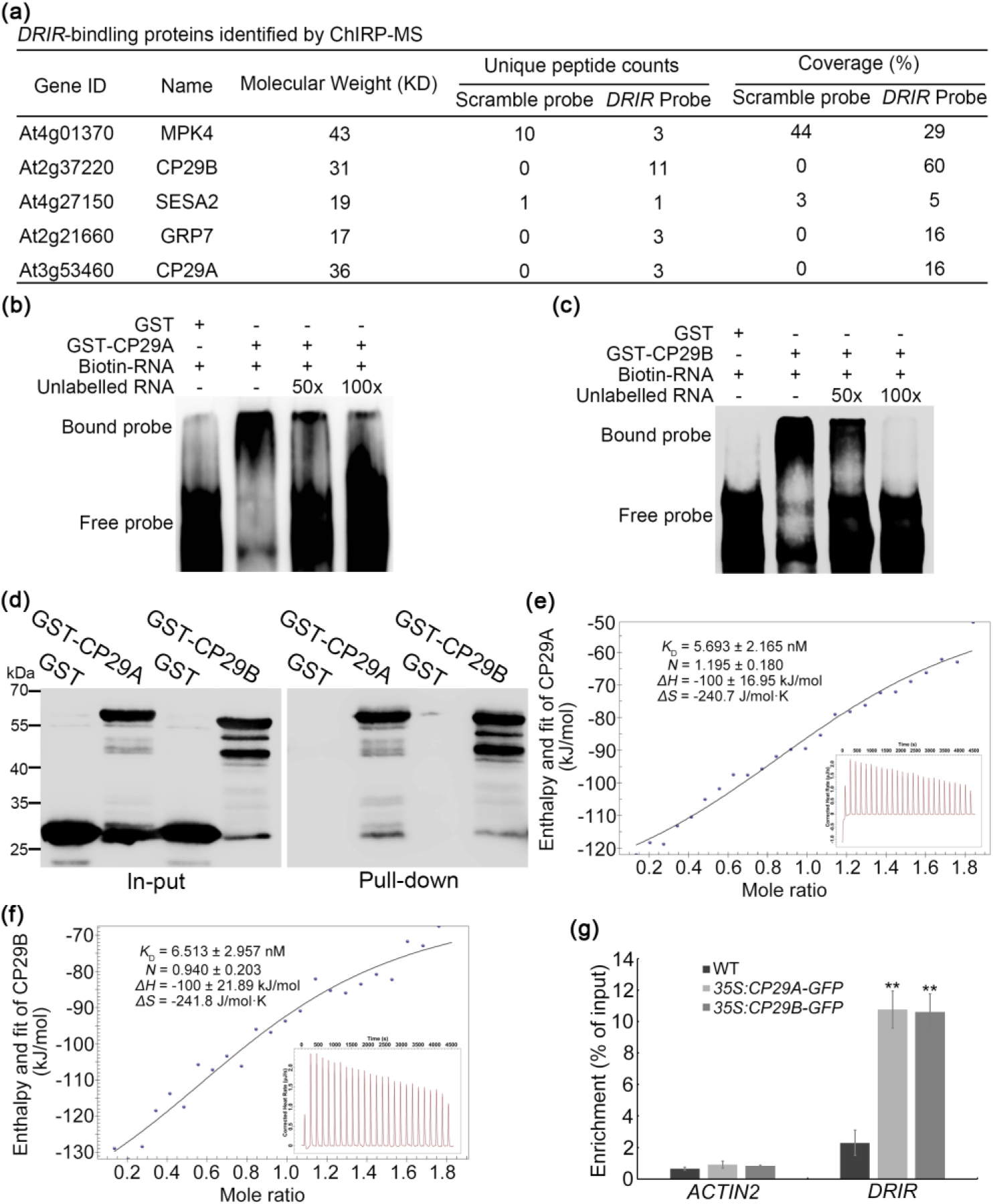
D*R*IR binds to CP29A and CP29B. (a) Potential *DRIR*-binding proteins identified by ChIRP-MS. A scramble probe served as a negative control. (b and c) RNA-EMSA showing the binding of *DRIR* to GST-CP29A (b) and GST-CP29B (c). The GST tag alone served as a negative control. (d) RNA pull-down assay showing *DRIR* binding to GST-CP29A and GST-CP29B. The GST tag alone was used as a negative control. (e and f) Binding of *DRIR* to GST-CP29A (e) and GST-CP29B (f) in ITC assays normalized against the GST tag alone. K_D_, dissociation constant; N, stoichiometry; ΔH, enthalpy; ΔS, entropy. (g) RIP-qPCR showing the binding of CP29A and CP29B to the *DRIR* transcript in vivo. *ACTIN2* mRNA served as a negative control. Data represent means ± SD from three biological replicates. **, *P* < 0.01 by Student’s *t* test compared to WT.

### LncRNA *DRIR* negatively regulates plant tolerance to freezing stress

Our previous study showed that *DRIR* positively regulates plant tolerance to drought stress (Qin *et al*., 2017). When *cp29a* and *cp29b* were treated with drought stress, both showed improved growth and better survival compared with the wild-type (WT) (Fig. S2), in contrast to *DRIR*’s role in regulating drought tolerance. Since CP29A is required for efficient chloroplast RNA processing under low temperature conditions (Kupsch *et al*., 2012), we transferred 7-day-old seedlings of WT, *drir^D^* (a T-DNA insertion mutant with a higher *DRIR* expression), *DRIR* overexpressing (OE), *cp29a*, and *cp29b* grown under normal conditions to a 4°C growth chamber for an additional one-month growth period. Unexpectedly, only the central rosette leaves of *cp29a* exhibited bleaching; the coloration of young tissues in *drir^D^*, *DRIR* OE, and *cp29b* was indistinguishable from that of the WT (Fig. S3), suggesting that the primary functions of *DRIR* and CP29B may not be directly related to chloroplast activity during cold stress. Intriguingly, we found that *DRIR* regulates plant freezing tolerance. When 6-day-old WT seedlings were exposed to -3°C freezing treatment for 1 h, the quantitative real-time PCR (qRT-PCR) results showed that the expression level of *DRIR* was significantly downregulated to approximately one-third of its level under normal conditions (22°C) (Fig. 2a). Similarly, when P*_DRIR_:GUS* transgenic seedlings were subjected to the same treatment, the β-Glucuronidase (GUS) signal intensity was significantly reduced compared to that under normal conditions (Fig. 2b), demonstrating that *DRIR* expression is responsive to freezing stress. When 14-day-old WT, *drir^D^*, and *DRIR* OE seedlings were subjected to -4°C treatment for 1 h, the survival rates of *drir^D^*, OE#1, and OE#2 were significantly lower than that of WT (Fig. 2c,d), being 0.61, 0.27, and 0.20 times that of WT, respectively (Fig. 2e). Correspondingly, the extent of plasma membrane injury in *drir^D^*, OE #1, and OE #2 was higher than that in WT, with electrolyte leakage being 1.11, 1.29, and 1.22 times that of WT, respectively (Fig. 2f). When seedlings were acclimated at 4°C for 4 days and then subjected to freezing stress at -6°C for 0.5 h, the survival rates of OE#1 and OE#2 seedlings were also significantly lower than that of WT. Meanwhile, their electrolyte leakage was 1.69 and 1.57 times that of WT, respectively (Fig. 2d-f). the observation that the freezing tolerance of *drir^D^* seedlings was higher than that of OE#1 and OE#2 is consistent with the significantly lower expression level of *DRIR* in *drir^D^* compared to OE#1 and OE#2 (Fig. 2c). To further elucidate the function of *DRIR* in plant freezing tolerance, we generated *DRIR* RNAi knockdown lines (Fig. 2g). These lines displayed significantly improved freezing tolerance (Fig. 2h). Compared to WT, the survival rates of *DRIR* RNAi seedlings increased by over 44% and 65% under non-acclimated and cold-acclimated conditions, respectively. Correspondingly, electrolyte leakage decreased by over 19% and 25% (Fig. 2i,j). These experimental results indicate that the expression level of lncRNA *DRIR* influences the freezing tolerance of plant seedlings.

**Fig. 2.**
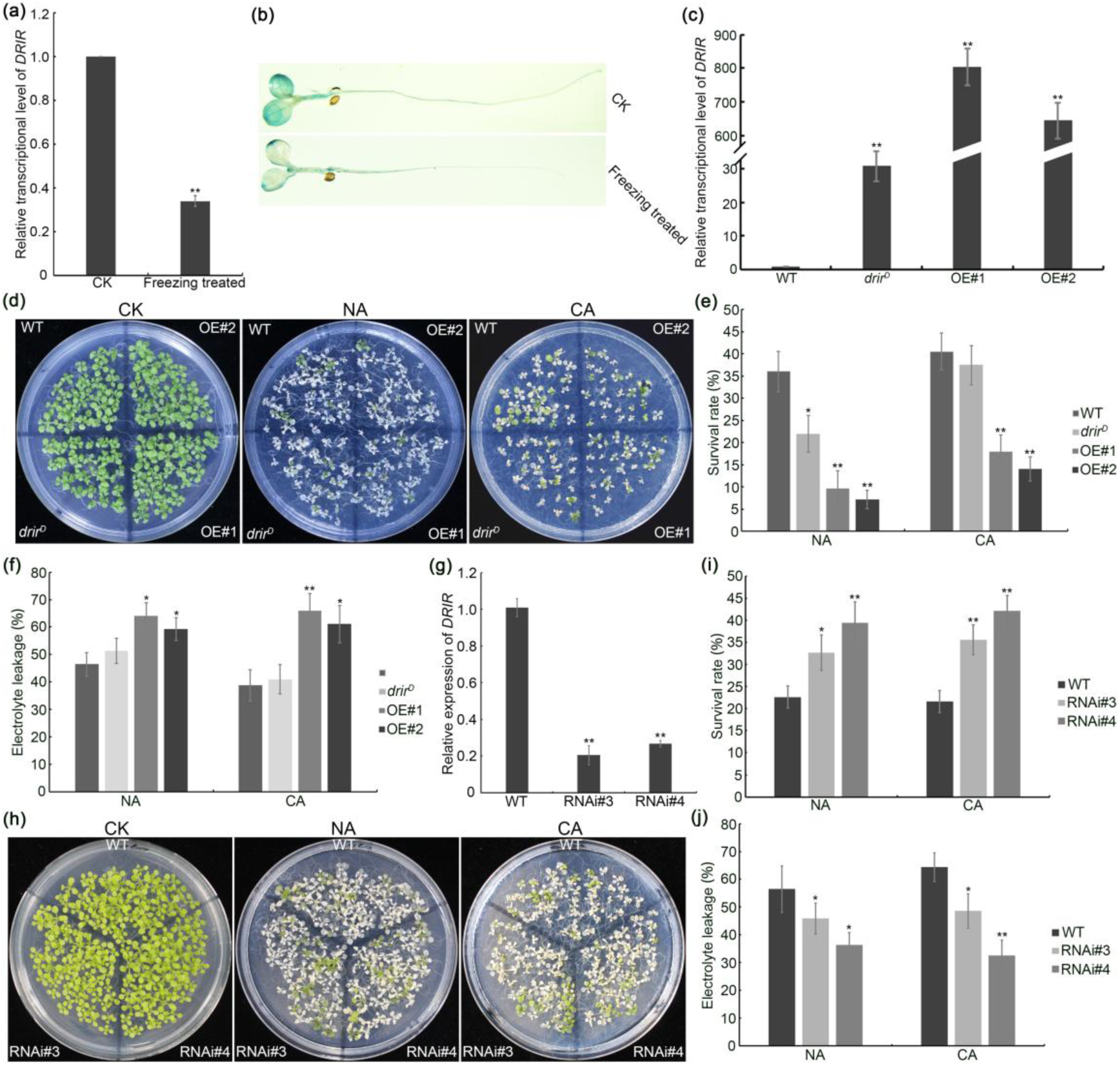
D*R*IR regulates freezing tolerance in plant seedlings. (a) The expression of *DRIR* in seedlings subjected to -3°C freezing treatment for 1 h. Data represent means ± SD from three biological replicates. (b) GUS signal of *P_DRIR_:GUS* transgenic seedlings with or without freezing treatment. (c) Expression levels of *DRIR* in wild-type (WT), *drir^D^*, and two *DRIR* overexpressing (OE) lines. Data represent means ± SD from three biological replicates. (d-f) Representative images (d), survival rates (e), and electrolyte leakage (f) of WT, *drir^D^*, and OE seedlings under non-acclimated (NA) and acclimated (CA; 4 days at 4°C) conditions. Two-week-old seedlings were subjected to -4°C for 1 h (NA) or -6°C for 0.5 h (CA). Data represent means ± SD (n > 10 plates for survival rates; n = 3 biological replicates for electrolyte leakage). (g) Expression levels of *DRIR* in two representative *DRIR* RNAi lines. Data represent means ± SD from three biological replicates. (h-j) Representative images (h), survival rates (i), and electrolyte leakage (j) of WT and *DRIR* RNAi seedlings under non-acclimated and acclimated conditions. Data represent means ± SD (n > 10 plates for survival rate and n = 3 biological replicates for electrolyte leakage). Statistical significance was determined using the Student’s *t* test. * *P* < 0.05; ** *P* < 0.01.

### CP29A and CP29B also affect plant tolerance to freezing stress

To investigate the involvement of CP29A and CP29B in plant tolerance to freezing stress, we exposed *cp29a* and *cp29b* to -4°C freezing stress. The results showed that both mutants exhibited lower survival rates and higher electrolyte leakage compared to the WT (Fig. 3a-c). When seedlings were cold-acclimated at 4°C for 4 days followed by -6°C freezing treatment for 0.5 h, the survival rates of *cp29a* and *cp29b* seedlings were 55% and 31% of the WT survival rate, respectively. Correspondingly, their electrolyte leakage was 1.56- and 1.89-fold that of the WT, respectively (Fig. 3a-c). These results indicate that *cp29a* and *cp29b* are more susceptible to freezing stress. We also generated CP29A- and CP29B-overexpressing (OE) transgenic lines and subjected them to freezing stress treatment. The results demonstrated that CP29A-OE and CP29B-OE seedlings exhibited enhanced freezing tolerance compared with the WT regardless of cold acclimation status. This was evidenced by significantly higher survival rates and lower electrolyte leakage (Fig. 3d-i). Collectively, these phenotypic data indicate that CP29A and CP29B can affect plant tolerance to freezing stress.

**Fig. 3.**
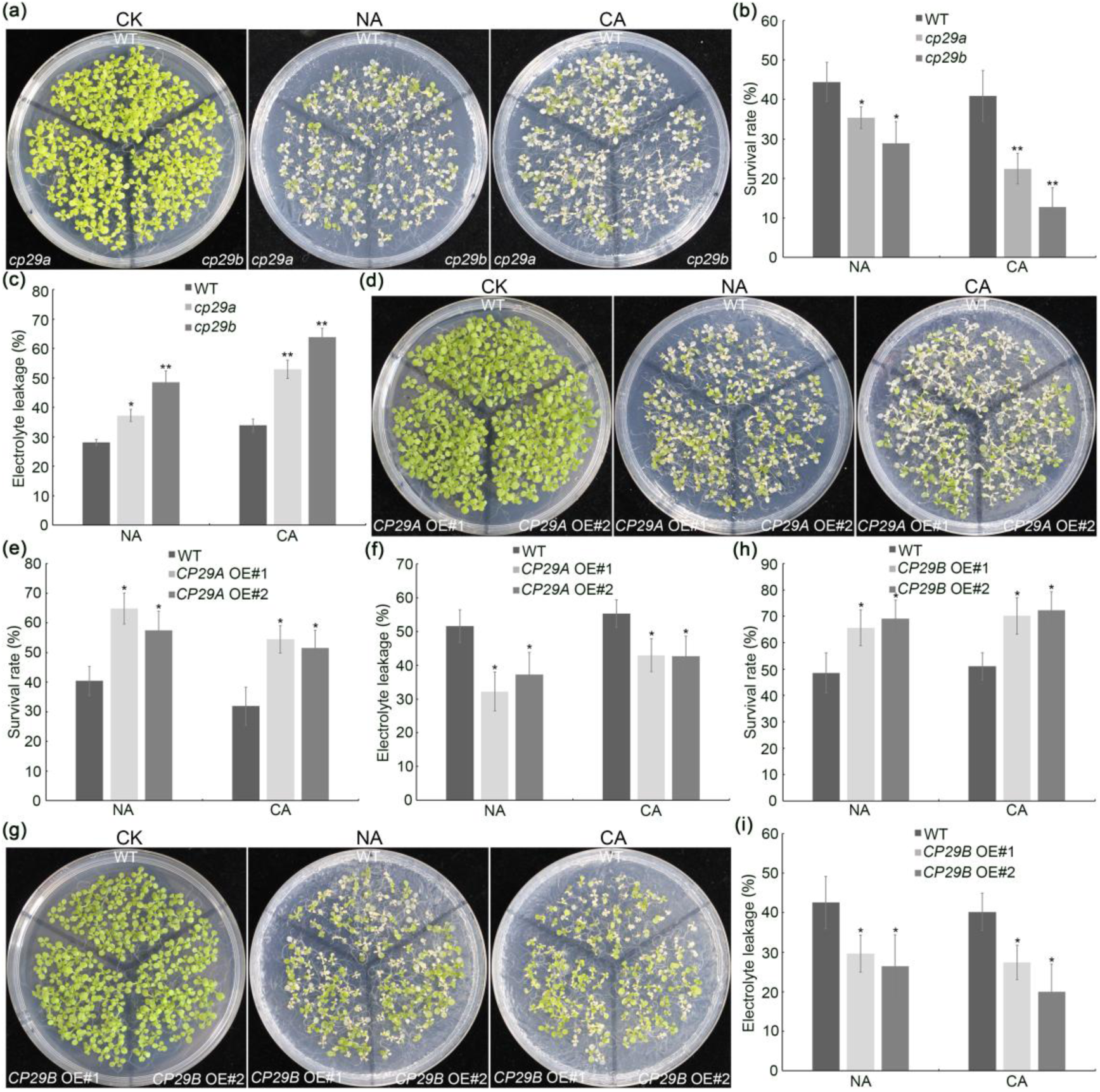
CP29A and CP29B affect tolerance to freezing stress. (a-c) Representative images (a), survival rates (b), and electrolyte leakage (c) of WT, *cp29a*, and *cp29b* seedlings under non-acclimated and acclimated conditions. Data represent means ± SD (n > 10 plates for survival rates; n = 3 biological replicates for electrolyte leakage). (d-f) Representative images (d), survival rates (e), and electrolyte leakage (f) of WT and *CP29A* OE seedlings under non-acclimated and acclimated conditions. Data represent means ± SD (n > 10 plates for survival rates; n = 3 biological replicates for electrolyte leakage). (g-i) Representative images (g), survival rates (h), and electrolyte leakage (i) of WT and *CP29B* OE seedlings under non- acclimated and acclimated conditions. Data represent means ± SD (n > 10 plates for survival rates; n = 3 biological replicates for electrolyte leakage). Statistical significance was determined using the Student’s *t* test. * *P* < 0.05; ** *P* < 0.01.

### *DRIR*-CP29A/CP29B module affects mRNA alternative splicing

Previous research indicated that cpRNPs mainly regulate RNA maturation, pre-mRNA splicing, and RNA editing (Nakamura *et al*., 2001; Teubner *et al*., 2017). To investigate whether *DRIR* regulates RNA processing, we analyzed transcriptome data from our previous study on *drir^D^* and *DRIR* OE under drought stress (Qin *et al*., 2017) and found that *DRIR* influenced alternative splicing of pre-mRNA. Compared to the WT, *drir^D^* and *DRIR* OE shared 446 overlapping differential splicing events (Fig. S4a). Among these events, a significant downregulation of retained intron (RI) and skipped exon (SE) were observed in *drir^D^* and *DRIR* OE. Additionally, alternative 5’ splice site (A5’SS) and alternative 3’ splice site (A3’SS) events also differed from those in WT (Fig. S4a). RT-PCR validation of differential splicing events showed consistent results with the transcriptome sequencing data. In *drir^D^* and *DRIR* OE, the relative splicing levels of representative RI (At2g25850), SE (At3g05820), A5’SS (At3g27610), and A3’SS (At2g25850) were lower than in WT (Fig. S4b). To explore whether the *DRIR*-CP29A/CP29B module regulates plant freezing resistance by affecting alternative splicing of pre-mRNA, we performed transcriptome analysis on WT, *drir^D^*, *DRIR* OE, *cp29a*, and *cp29b* seedlings after freezing treatment. The results showed that *drir^D^*, *DRIR* OE, *cp29a*, and *cp29b* had 1644, 1642, 1553, and 1611 differential splicing events compared to WT, with 1025, 1155, 977, and 1083 of these events being downregulated, respectively (Fig. 4a). Analysis of these splicing events revealed that *drir^D^*, *DRIR* OE, *cp29a*, and *cp29b* shared 74 upregulated and 285 downregulated differential splicing events (Fig. 4b). However, only 10 differential splicing events showed opposite regulation -- they were upregulated in *drir^D^* and *DRIR* OE but downregulated in *cp29a* and *cp29b*, and vice versa (Fig. S5). This suggests a positive correlation between the effects of *drir^D^* and *DRIR* OE on pre-mRNA splicing and those of *cp29a* and *cp29b*. Analysis of the types of differential splicing events revealed that the numbers of downregulated RI, SE, mutually exclusive exon (MXE), A5’SS, and A3’SS events in *drir^D^*, *DRIR* OE, *cp29a*, and *cp29b* were significantly greater than the numbers of upregulated events (Fig. 4c). Validation of representative differential splicing events confirmed that, consistent with the transcriptome sequencing results, the splicing ratios (the level of alternative splicing isoform divided by that of constitutive splicing isoform) of RI for At2g17840, At5g04980, and At1g62660 in *drir^D^*, *DRIR* OE, *cp29a*, and *cp29b* seedlings were lower than those in WT (Fig. 4d). Similarly, the splicing ratios for SE (At5g51820), A5’SS (At4g01870), and A3’SS (At1g23230) were also lower than in WT (Fig. 4e-g). To confirm that CP29A and CP29B can act on nuclear transcripts, we performed RIP-qPCR to verify their specific binding to these target RNAs, even though nuclear transcript association was previously reported (Gosai *et al*., 2015). The results demonstrated that all these target transcripts were significantly enriched in the immunoprecipitates of *35S:CP29A-GFP* and *35S:CP29B-GFP* (Fig. 4h). These results indicate that the *DRIR*-CP29A/CP29B module regulates pre-mRNA splicing under freezing stress.

**Fig. 4.**
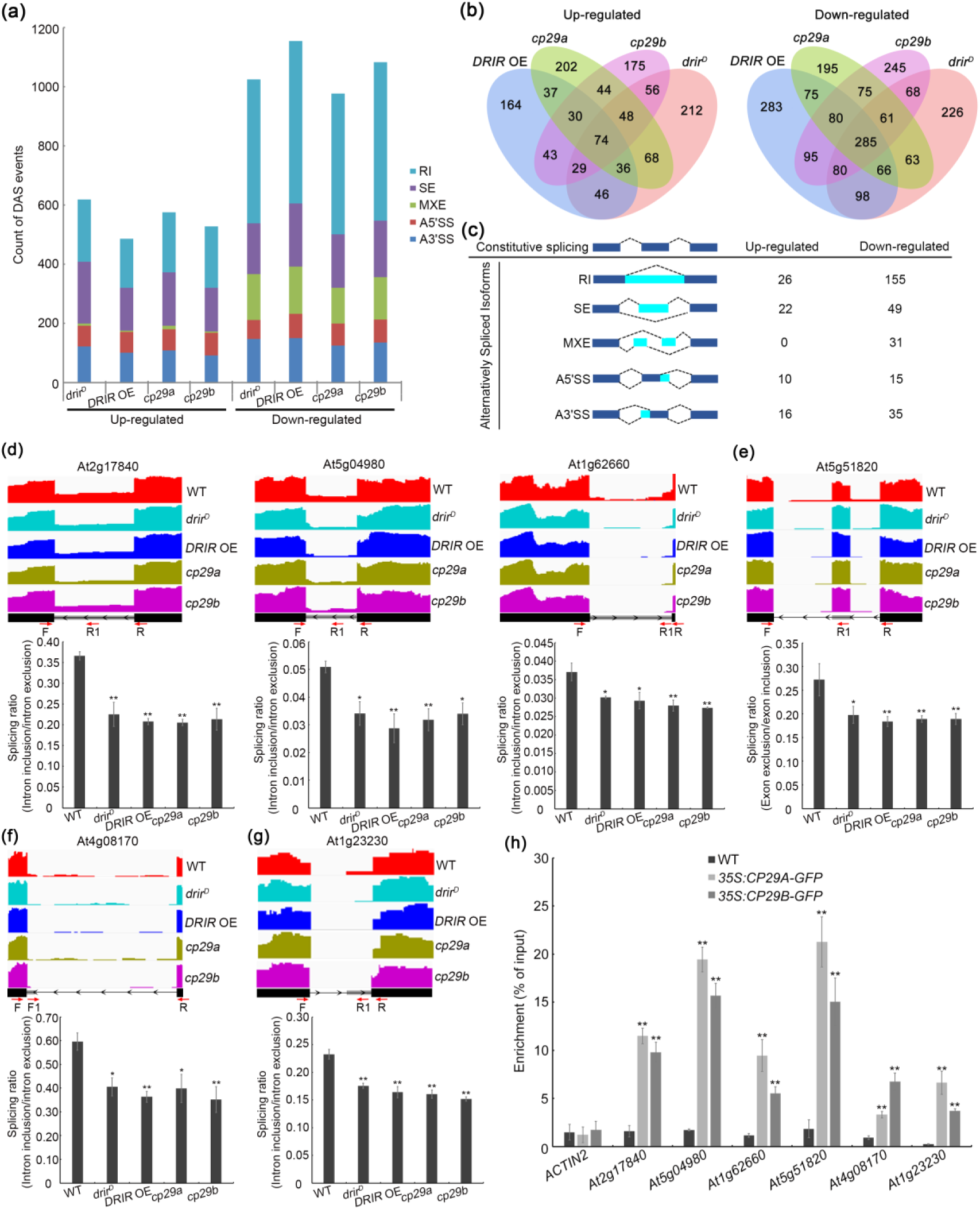
Effects of *DRIR*, CP29A, and CP29B on pre-mRNA alternative splicing under freezing stress. (a) Count of differential alternative splicing (DAS) events in *drir^D^*, *DRIR* OE, *cp29a*, and *cp29b* seedlings compared to WT. Six-day-old seedlings were subjected to -3°C freezing treatment for 1 h followed by transcriptome analysis. RI, retained intron; SE, skipped exon; MXE, mutually exclusive exon; A5’SS, alternative 5’ splice site; A3’SS, alternative 3’ splice site. (b) Venn diagrams showing the overlaps among the upregulated or downregulated differential alternative splicing events in *drir^D^*, *DRIR* OE, *cp29a*, and *cp29b*. (c) Types of co-upregulated or co-downregulated differential alternative splicing events in *drir^D^*, *DRIR* OE, *cp29a*, and *cp29b* seedlings. (d-g) Integrative Genomics Viewer (IGV) tracks and splicing ratios of representative alternative splicing events of RI (d), SE (e), A5’SS (f), and A3’SS (g) events in WT, *drir^D^*, *DRIR* OE, *cp29a*, and *cp29b* seedlings. IGV gene tracks display sequencing signal intensities for representative alternative splicing events in the transcriptomes. The exon-intron structure diagrams are shown below each IGV track. Black boxes define the exonic regions, introns are represented as lines and the alternative regions are indicated by gray boxes. Red arrows indicate the positions of the designed primers. Splicing efficiency is calculated as the ratio of intron inclusion to intron exclusion or exon exclusion to exon inclusion measured by RT-qPCR. Data represent means ± SD from three biological replicates. (h) RIP-qPCR assay showing CP29A and CP29B binding to their target transcripts. *ACTIN2* mRNA served as a negative control. Data represent means ± SD from three biological replicates. Statistical significance was determined using the Student’s *t* test. * *P* < 0.05; ** *P* < 0.01.

### *DRIR* promotes autophagy-mediated degradation of CP29A and CP29B

The *DRIR* gene lacks introns, eliminating pre-mRNA splicing possibilities. Therefore, CP29A and CP29B cannot function via regulating the splicing of *DRIR* RNA. To explore whether CP29A and CP29B affect the expression of *DRIR*, the transcript levels of *DRIR* in WT, *cp29a*, and *cp29b* seedlings were analyzed. It was found that the expression levels of *DRIR* in *cp29a* and *cp29b* were similar as that in WT (Fig. S6), suggesting that *DRIR* expression is unlikely to be controlled by CP29A and CP29B. Since lncRNAs can regulate the localization and activity of their interacting proteins (Kornienko *et al*., 2024), we first examined whether *DRIR* affects the subcellular localization of CP29A and CP29B. When we transiently expressed *35S:CP29A-GFP* or *35S:CP29B-GFP* with or without *35S:DRIR* in tobacco leaves, the results showed that CP29A and CP29B were mainly localized in chloroplasts when *DRIR* was not co-expressed. In contrast, numerous non-chloroplast granular signals of CP29A and CP29B were observed when co-expressed with *DRIR* (Fig. 5a). Next, we transformed *35S:CP29A-GFP* and *35S:CP29B-GFP* into WT, *drir^D^*, and *DRIR* OE Arabidopsis protoplasts. Similarly, CP29A and CP29B were primarily localized in chloroplasts in WT protoplasts. However, in *drir^D^* and *DRIR* OE protoplasts, strong granular GFP signals appeared outside the chloroplasts, accompanied by weakened signals inside chloroplasts (Fig. 5b).

**Fig. 5.**
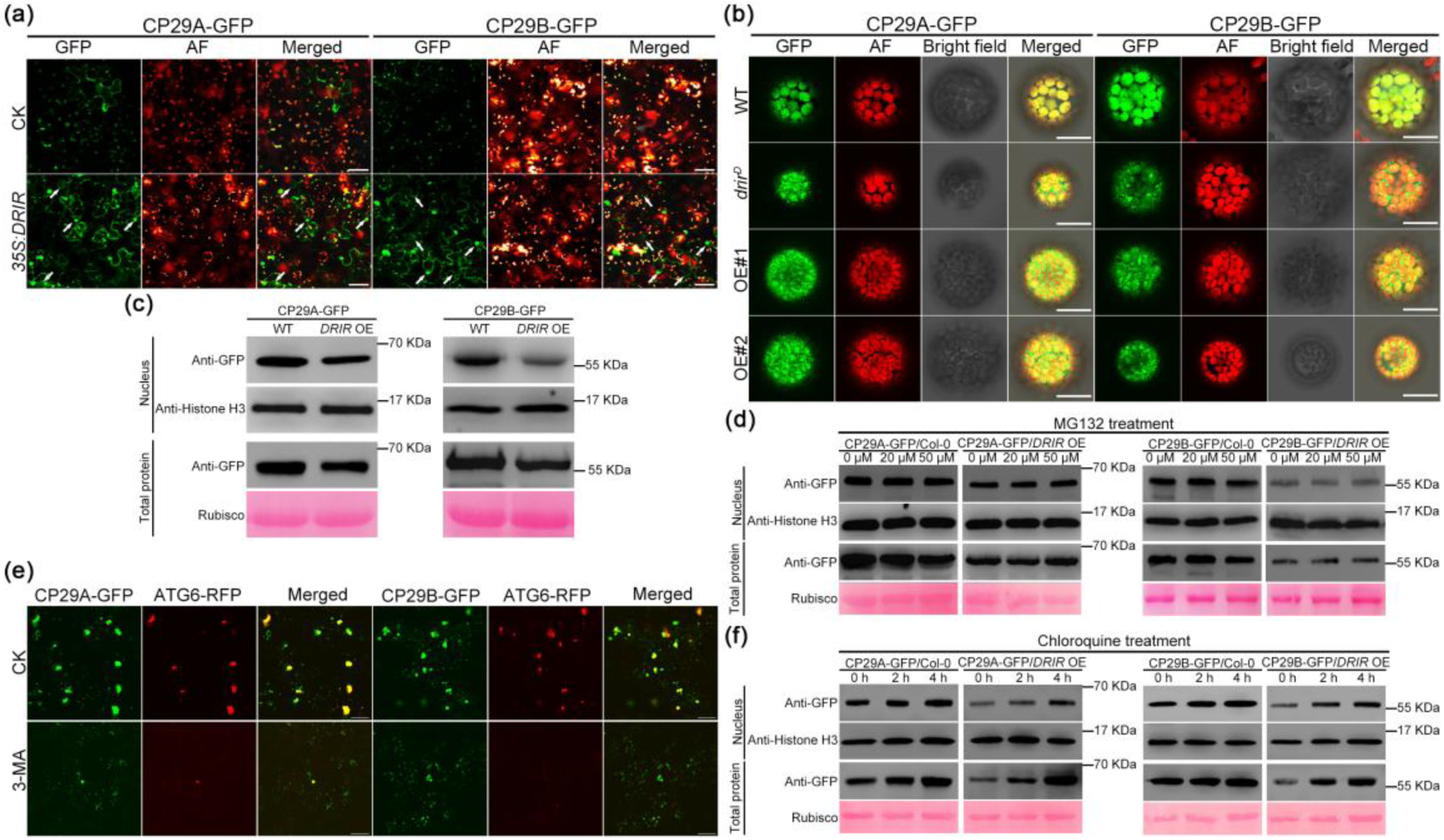
D*R*IR induces autophagy-based degradation of CP29A and CP29B. (a) Subcellular localization of CP29A and CP29B in tobacco leaves co-infiltrated with or without *35S:DRIR*. AF, chloroplast autofluorescence. Arrows indicate non-chloroplast granules. Scale bars = 50 μm. (b) Subcellular localization of CP29A and CP29B in protoplasts of WT, *drir^D^*, and *DRIR* OE. Scale bars = 20 μm. (c) Protein levels of CP29A and CP29B in total and nuclear protein fractions from WT and *DRIR* OE seedlings. Histone H3 and Rubisco served as immunoblot loading controls for nuclear and total proteins, respectively. (d) Protein levels of CP29A and CP29B in WT and *DRIR* OE after MG132 treatment. Histone H3 and Rubisco were used as immunoblot loading controls for nuclear and total proteins, respectively. (e) Co-localization of non-chloroplast granules of CP29A and CP29B with the autophagosome marker ATG6-RFP. Tabaco leaves were pre-treated with 0.1% DMSO (CK) or 10 mM 3-Methyladenine (3-MA) for 12 h. Scale bars = 50 μm. (f) Protein levels of CP29A and CP29B in WT and *DRIR* OE after chloroquine treatment. Histone H3 and Rubisco were used as immunoblot loading controls for nuclear and total proteins, respectively.

To explore whether the altered cellular localization of CP29A and CP29B affects their function in the nucleus, we extracted the total and nuclear proteins of WT and *DRIR* OE and detected the protein levels of CP29A and CP29B using western blotting. The results indicated that both total and nuclear protein levels of CP29A and CP29B in *DRIR* OE were reduced compared to WT (Fig. 5c). However, the *DRIR*-induced reduction of CP29A and CP29B was not prevented by MG132, indicating that CP29A and CP29B were not degraded in *DRIR* OE seedlings through the proteasome pathway (Fig. 5d). Therefore, we wondered whether it was regulated by another major degradation system: the autophagy pathway. To test the possibility of the involvement of autophagy, we investigated the co-localization of the granular signals of CP29A and CP29B with autophagosomes. The results showed that non-chloroplast granular signals of CP29A and CP29B in tobacco leaves coexpressing *35S:DRIR* were co-localized with the autophagosome marker ATG6-RFP. When treated with the specific inhibitor of autophagosome formation, 3-Methyladenine (3-MA) (Galluzzi *et al*., 2017), these granular signals were almost completely eliminated (Fig. 5e), indicating that *DRIR* could induce the relocalization of CP29A and CP29B to autophagosomes. Furthermore, when WT and *DRIR* OE seedlings were treated with 200 µM chloroquine to block autophagosome-lysosome fusion (Galluzzi *et al*., 2017), the protein levels of CP29A and CP29B in both nuclear and total protein fractions, particularly in *DRIR* OE, increased substantially (Fig. 5f). These data demonstrated that *DRIR* promotes the autophagosome-mediated degradation of CP29A and CP29B.

### Disruption of CP29A and CP29B attenuates the resistance of *DRIR* RNAi seedlings to freezing stress

To analyze the genetic relationship between *DRIR* and CP29A or CP29B in regulating freezing tolerance, we generated *DRIR* RNAi *cp29a* and *DRIR* RNAi *cp29b* double mutants through crossing and subjected their seedlings to freezing stress. The results showed that *cp29a* and *cp29b* mutations greatly compromised the freezing-tolerant phenotype of *DRIR* RNAi. When subjected to -4°C treatment for 1 h, the survival rates of WT, *cp29a*, *DRIR* RNAi *cp29a* double mutant, and *DRIR* RNAi were 30%, 18%, 28%, and 56%, respectively. Meanwhile, their electrolyte leakage was 52%, 68%, 64%, and 40%, respectively (Fig. 6a-c). Similarly, *DRIR* RNAi *cp29a* seedlings exhibited only 47% of the WT survival rate and had 1.27-fold higher electrolyte leakage than WT after -6°C treatment for 0.5 h following cold acclimation (Fig. 6a-c). Similar trends were observed in the *DRIR* RNAi *cp29b* mutant. The survival rates of *DRIR* RNAi *cp29b* seedlings were 60% and 74% of WT levels under non-acclimated and cold-acclimated conditions, respectively (Fig. 6d,e). Correspondingly, the electrolyte leakage in *DRIR* RNAi *cp29b* mutant was 1.34 and 1.28 times that of WT, respectively (Fig. 6f). These results demonstrate that *CP29A* and *CP29B* act as downstream functional genes in *DRIR*-regulated freezing tolerance in plant seedlings.

**Fig. 6.**
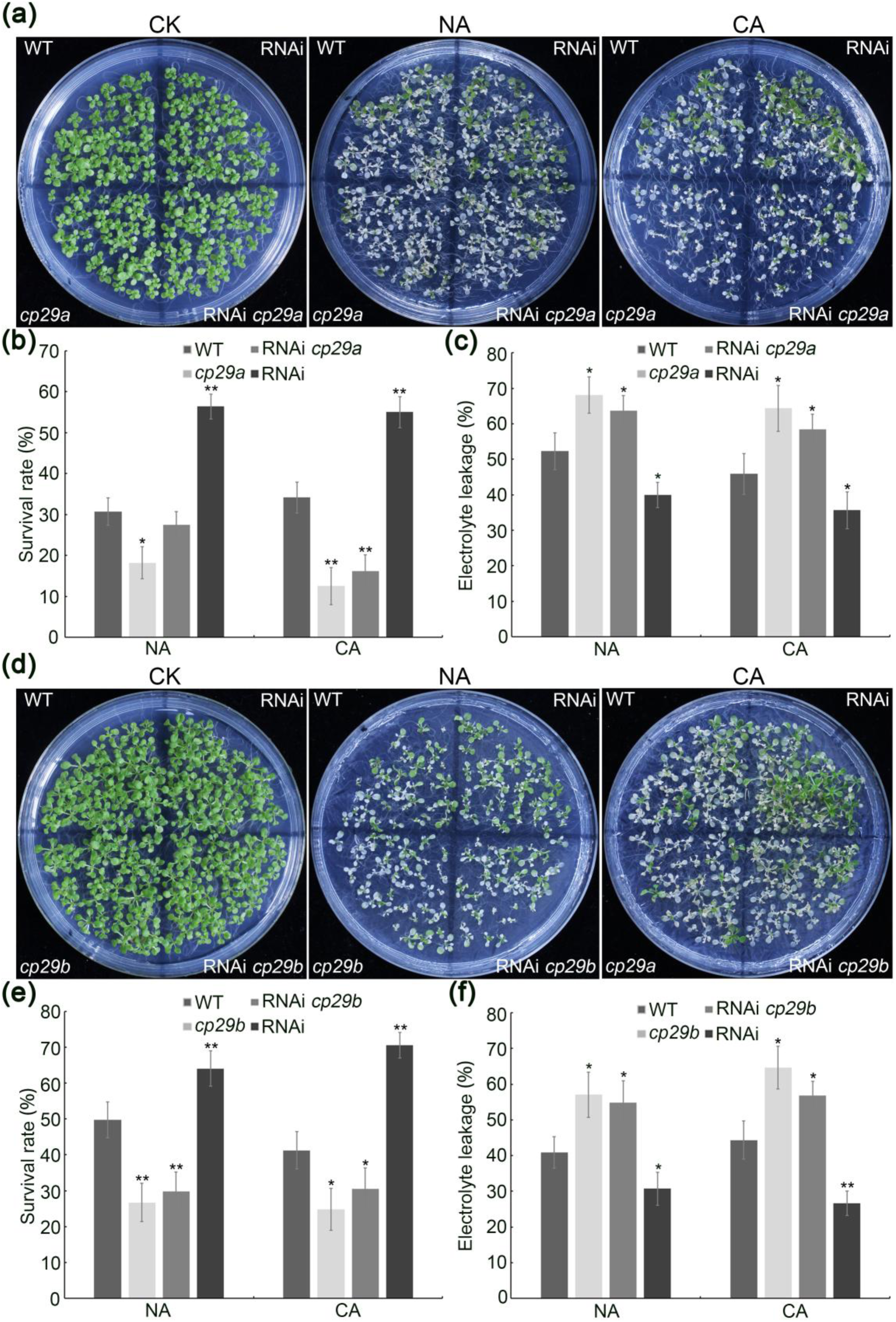
c*p*29a and *cp29b* compromise the freezing-tolerant phenotype of *DRIR* RNAi. (a-c) Representative images (a), survival rates (b), and electrolyte leakage (c) of WT, *cp29a*, *DRIR* RNAi *cp29a* double mutant, and *DRIR* RNAi seedlings under non-acclimated and acclimated conditions. Data represent means ± SD (n > 10 plates for survival rates; n = 3 biological replicates for electrolyte leakage). (d-f) Representative images (d), survival rates (e), and electrolyte leakage (f) of WT, *cp29b*, *DRIR* RNAi *cp29b* double mutant, and *DRIR* RNAi seedlings. Data represent means ± SD (n > 10 plates for survival rates; n = 3 biological replicates for electrolyte leakage). Statistical significance was determined using the Student’s *t* test. * *P* < 0.05; ** *P* < 0.01.

## Discussion

There is increasing evidence that lncRNAs are essential regulators of a range of biological processes (Yu *et al*., 2019; Gonzales *et al*., 2024). However, understanding the intricate molecular mechanisms of stress-responsive lncRNAs in plants is in its infancy. Even in the model plant *Arabidopsis thaliana*, only lncRNAs of the *SVALKA*-*asCBF1* cascade, *asZAT5*, *asBBX28*, and *CIL1* involved in regulating plant freezing tolerance have been functionally described—a number that lags far behind the quantity of lncRNAs identified in response to low temperatures (Kindgren *et al*., 2018; Liu *et al*., 2022; Meena *et al*., 2024). Here, we identified that the lncRNA *DRIR* plays a role in plant response to freezing stress. The seedlings of the *drir^D^* mutant and OE lines showed increased susceptibility to freezing stress, while *DRIR* RNAi lines exhibited enhanced tolerance compared to WT. Under non-cold-acclimated conditions, the survival rates of *drir^D^* and OE seedlings were significantly lower than that of WT. After cold acclimation, although the freezing tolerance of all genotypes improved (as evidenced by the survival rates and electrolyte leakage under more severe stress conditions at -6°C), the survival rates of *drir^D^* and OE seedlings remained lower than that of WT (Fig. 2).

Low-temperature stress induces massive and rapid alternative splicing alteration coincident with changes in gene transcription levels (Calixto *et al*., 2018). Differential splicing of certain genes plays a significant role in the cold stress response, including freezing tolerance (Seo *et al*., 2012; Guan *et al*., 2013; Wang *et al*., 2022; Zhong *et al*., 2024). REGULATOR OF CBF GENE EXPRESSION1 (RCF1) ensures the proper pre-mRNA splicing of many genes under cold stress, leading to the plant’s tolerance to both chilling and freezing stress (Guan *et al*., 2013). In contrast, Sm protein E1 (SME1), a core component of the spliceosome, is essential for the efficiency of constitutive and alternative splicing of pre-mRNAs and negatively regulates plant tolerance to freezing stress (Huertas *et al*., 2019). lncRNAs have emerged as regulators of alternative splicing through various molecular mechanisms (Gonzales *et al*., 2024). In the model legume *Medicago truncatula*, the lncRNA *EARLY NODULIN 40* (*ENOD40*) interacts with and regulates the nucleocytoplasmic trafficking of the RNA-binding protein 1 (RBP1), affecting RBP1-dependent alternative splicing events (Campalans *et al*., 2004). *ALTERNATIVE SPLICING COMPETITOR* (*ASCO*), the homolog of *ENOD40* in *Arabidopsis*, competes with alternatively spliced mRNA targets to bind nuclear speckle RNA-binding protein (NSR), thus affecting NSR-dependent alternative splicing events as well (Bardou *et al*.,2014). In addition to NSRs, *ASCO* also interacts with PRE-MRNA PROCESSING 8A (PRP8A) and Sm ring D1b (SmD1b), which are core components of the spliceosome, to regulate alternative splicing (Rigo *et al*., 2020). The *FLOWERING ASSOCIATED INTERGENIC lncRNA* (*FLAIL*) affects alternative splicing in *Arabidopsi*s by acting as a scaffold to bridge RNA-RNA, RNA-DNA, and RNA-protein interactions to facilitate the splicing of specific targets (Jin *et al*., 2023). A long circular RNA (*circRNA*) transcribed from the *SEPALLATA3* (*Sep3*) regulates splicing of its cognate mRNA by forming a DNA–RNA duplex R-loop (Conn *et al*., 2017). However, the role of lncRNA-mediated alternative splicing in plant response to low-temperature stress has not yet been elucidated.

In our previous study, we found that *DRIR* is a long intergenic ncRNA (lincRNA) transcribed from an intergenic region. It neither affects the expression of its neighboring genes nor acts as a precursor or target mimic for miRNA. Fluorescence in situ hybridization showed that *DRIR* is primarily localized in the nucleus and may regulate stress-responsive gene expression via an unknown nuclear mechanism (Qin *et al*., 2017). However, the specific molecular mechanism remains unclear. In this study, we found that *DRIR* affected alternative splicing in plant response to freezing stress. Using the ChIRP-MS method, we identified two potential interacting proteins of *DRIR*. Further studies, including EMSA, RNA-protein pull-down, ITC, and RIP-qPCR assays, confirmed that CP29A and CP29B can directly bind to *DRIR* both in vivo and in vitro (Fig. 1). Previous research has shown that CP29A is essential for efficient chloroplast RNA splicing, as RNA splicing of chloroplast introns is significantly reduced in the *cp29a* mutants (Kupsch *et al*., 2012; Legen *et al*., 2024). In addition to chloroplasts, CP29A also localizes to the nucleus and binds to nuclear mRNA containing GAN repeat motif, which is believed to be associated with alternative splicing (Wu *et al*., 2014; Gosai *et al*., 2015). Our results revealed that *DRIR*, along with CP29A and CP29B, can jointly influence alternative splicing of nuclear pre-mRNAs in plants. Transcriptome sequencing revealed that seedlings of *drir^D^*, *DRIR* OE, *cp29a*, and *cp29b* had more down-regulated differential splicing events. The splicing ratios (defined as the level of alternative splicing isoform relative to constitutive splicing isoform) of representative genes were lower in these genotypes than in WT (Fig. 4a-g, S4). Western blot analysis of total and nuclear protein fractions demonstrated that both CP29A and CP29B were identified in the nuclear protein fraction (Fig. 5c). RIP-qPCR results indicated that CP29A and CP29B directly bind to the splicing target transcripts (Fig. 4h). Rapid response to environmental changes is critical for plant adaptation to abiotic stress. Pre-mRNA splicing regulatory mechanisms, including alternative splicing, are among the important post-transcriptional mechanisms required for reprogramming gene expression and delivering regulatory plasticity in response to environmental stresses (Laloum *et al*., 2018; Huertas *et al*., 2019). In addition to affecting stress-responsive expression under drought conditions (Qin *et al*., 2017), our data showed that *DRIR* also influenced alternative splicing under these conditions, suggesting that *DRIR* possibly fine-tunes gene expression via the alternative splicing pathway. Disruption of the proper pre-mRNA splicing under cold stress may lead to altered plant tolerance to freezing stress (Guan *et al*., 2013; Huertas *et al*., 2019). Therefore, these data suggest that *DRIR* influences plant response to freezing stress via regulating pre-mRNA splicing, a mechanism that differs from other lncRNAs reported to respond to freezing stress (Kindgren *et al*., 2018; Liu *et al*., 2022; Meena *et al*., 2024).

When associated with proteins, lncRNAs usually function as molecular scaffolds to recruit target proteins to specific gene loci, or as decoys to sequester proteins from their targets (Zhao *et al*., 2018; Moison *et al*., 2021; Zhou *et al*., 2021; Zhu *et al*., 2021; Roule *et al*., 2022). Currently, the mechanisms by which plant lncRNAs interact with peripheral splicing factors or spliceosome core components to modulate alternative splicing mainly involve regulating the nucleocytoplasmic trafficking of the splicing factors, competing with target mRNAs for binding to splicing regulators, or mediating spliceosomal interactions with target mRNAs (Campalans *et al*., 2004; Bardou *et al*., 2014; Rigo *et al*., 2020; Lucero *et al*., 2021). However, we found *DRIR* regulates CP29A and CP29B through a mechanism not yet described in plants. In *drir^D^*or *DRIR* OE lines, CP29A and CP29B were abnormally relocated as granules outside the chloroplasts, and these granules co-localized with autophagosomes. Western blot assays revealed that the protein levels of CP29A and CP29B were reduced in both the total protein and nuclear fractions of the *DRIR* OE seedlings. MG132 treatment failed to restore their accumulation, while the reduction of CP29A and CP29B levels in *DRIR* OE could be inhibited by the autophagy inhibitor chloroquine (Fig. 5). These results indicate that *DRIR* can sequester CP29A and CP29B and mediate their autophagic degradation, thus interfering with alternative splicing.

Therefore, considering the massive and rapid alternative splicing alteration of pre-mRNAs contributes to stress plasticity under freezing conditions (Calixto *et al*., 2018; Huertas *et al*., 2019), our results uncover the following model for *DRIR* involved in plant response to freezing stress. Normally, freezing stress suppresses the expression levels of *DRIR*. Low levels of *DRIR* are unable to affect CP29A and CP29B in the nucleus by acting as a decoy. CP29A and CP29B enhance alternative splicing of pre-mRNAs under stress, thereby causing the rapid alteration of gene expression and improving plant freezing resistance. However, in the *drir^D^* and *DRIR* OE lines, elevated levels of *DRIR* bind to CP29A and CP29B in the nucleus, causing their relocalization to autophagosomes for degradation via the autophagic pathway. This, in turn, reduces the alternative splicing of nuclear RNAs, making the plants more sensitive to freezing stress (Fig. 7).

**Fig. 7.**
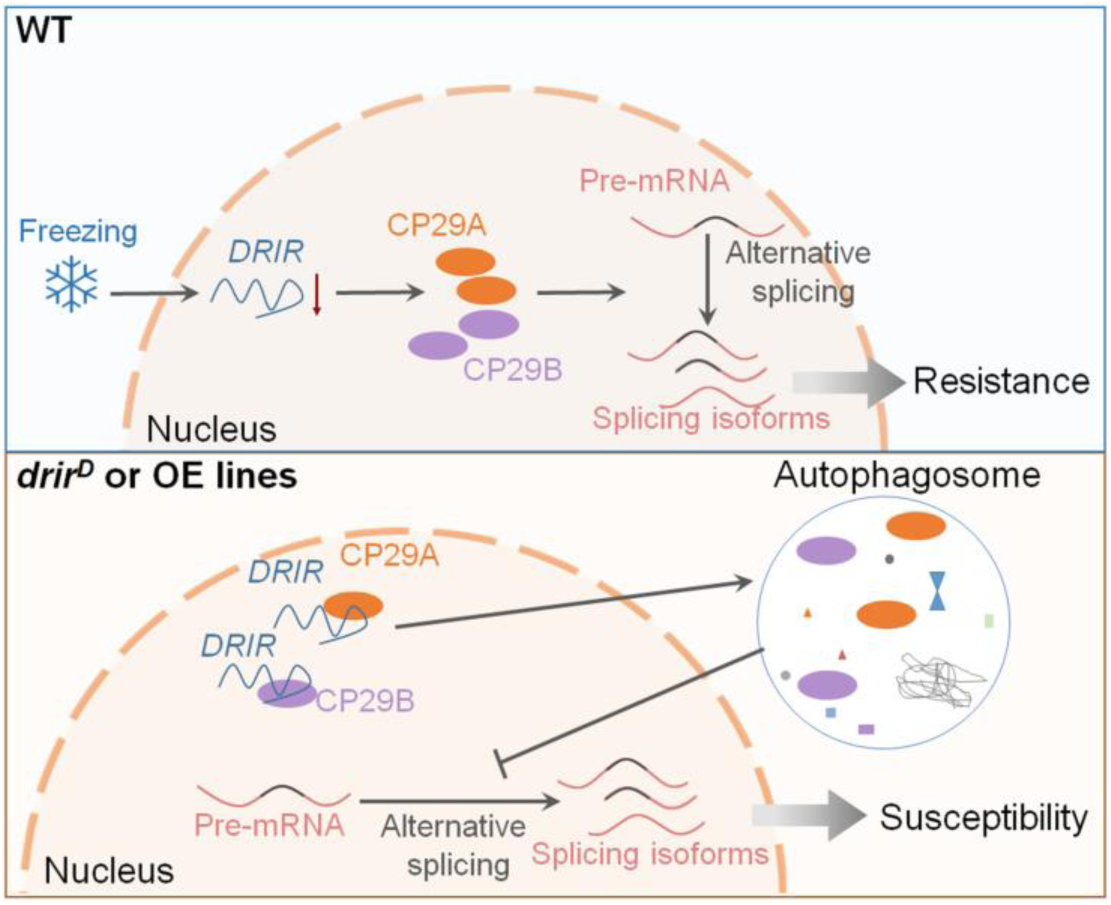
Model of *DRIR*’s role in plant freezing tolerance. In the wild-type plant, freezing stress reduces *DRIR* expression, limiting its ability to act as a decoy for CP29A and CP29B. This allows CP29A and CP29B to promote rapid alternative splicing of nuclear pre-mRNAs, enhancing plant freezing resistance. However, in *drir^D^* and *DRIR* OE seedlings, the elevated transcription level of *DRIR* leads to the relocalization of CP29A and CP29B into autophagosomes for degradation. Consequently, the normal stress-induced mRNA splicing reprogramming is disrupted, resulting in reduced plant freezing tolerance.

## Acknowledgments

This work was supported by the National Natural Science Foundation of China (32270315 to T.Q), Chinese Universities Scientific Fund (2452024381 to T.Q) and Research Grants Council (RGC) of Hong Kong General Research Fund (12101521 to L.X). We thank Dr. Qiuling Wang (Institute of Future Agriculture, Northwest A&F University) for providing the *35S:ATG6-RFP* plasmid; Dr. Fengping Yuan and Dr. Hua Zhao (State Key Laboratory for Crop Stress Resistance and High-Efficiency Production, Northwest A&F University, Yangling, China) for confocal experimental assistance.

## Author contributions

T.Q. and L.X. planned and designed the research. T.Q. supervised the work. L.Y., X.T., J.Y., and Z.Q. performed experiments or analyzed data. T.Q., C.W., and L.X. wrote the manuscript, revised and edited the article. All authors read and approved of its content.

## Data availability

The data that support the findings of this study are available in the supplementary material of this article (Figs S1–S6; Table S1). RNA-seq data from this article can be found in the Gene Expression Omnibus database (National Center for Biotechnology Information) under accession number GSE300716.

