## Supplementary Information for "The Function of LncRNA *DRIR* in Freezing Tolerance by Promoting Autophagic Degradation of CP29A and CP29B to Alter Alternative Splicing Patterns of Pre-mRNAs"

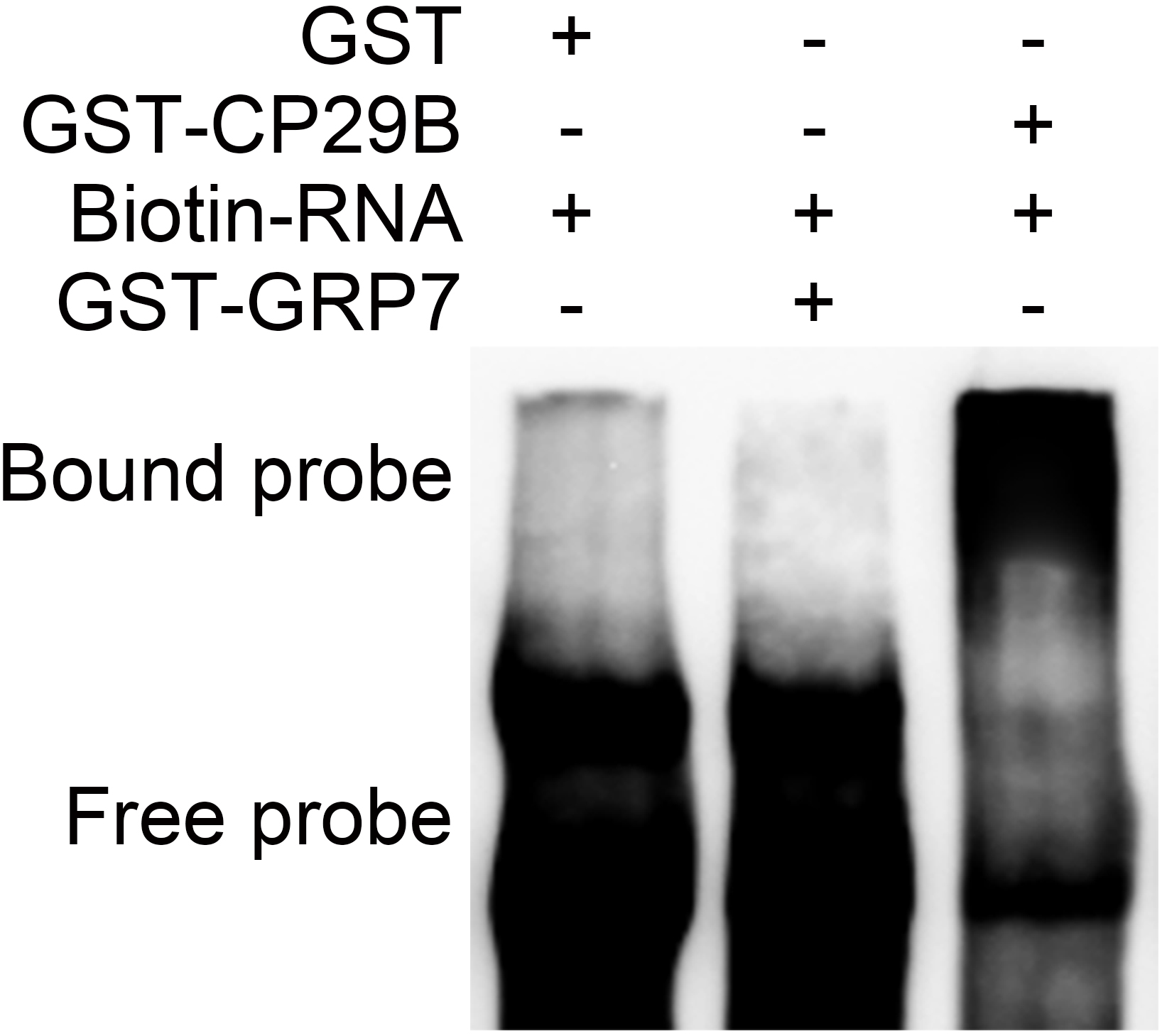

**Fig. S1** *DRIR* does not bind to GST-GRP7 in the RNA-EMSA assay. GST-CP29B and GST tag alone were used as the positive and negative control, respectively.

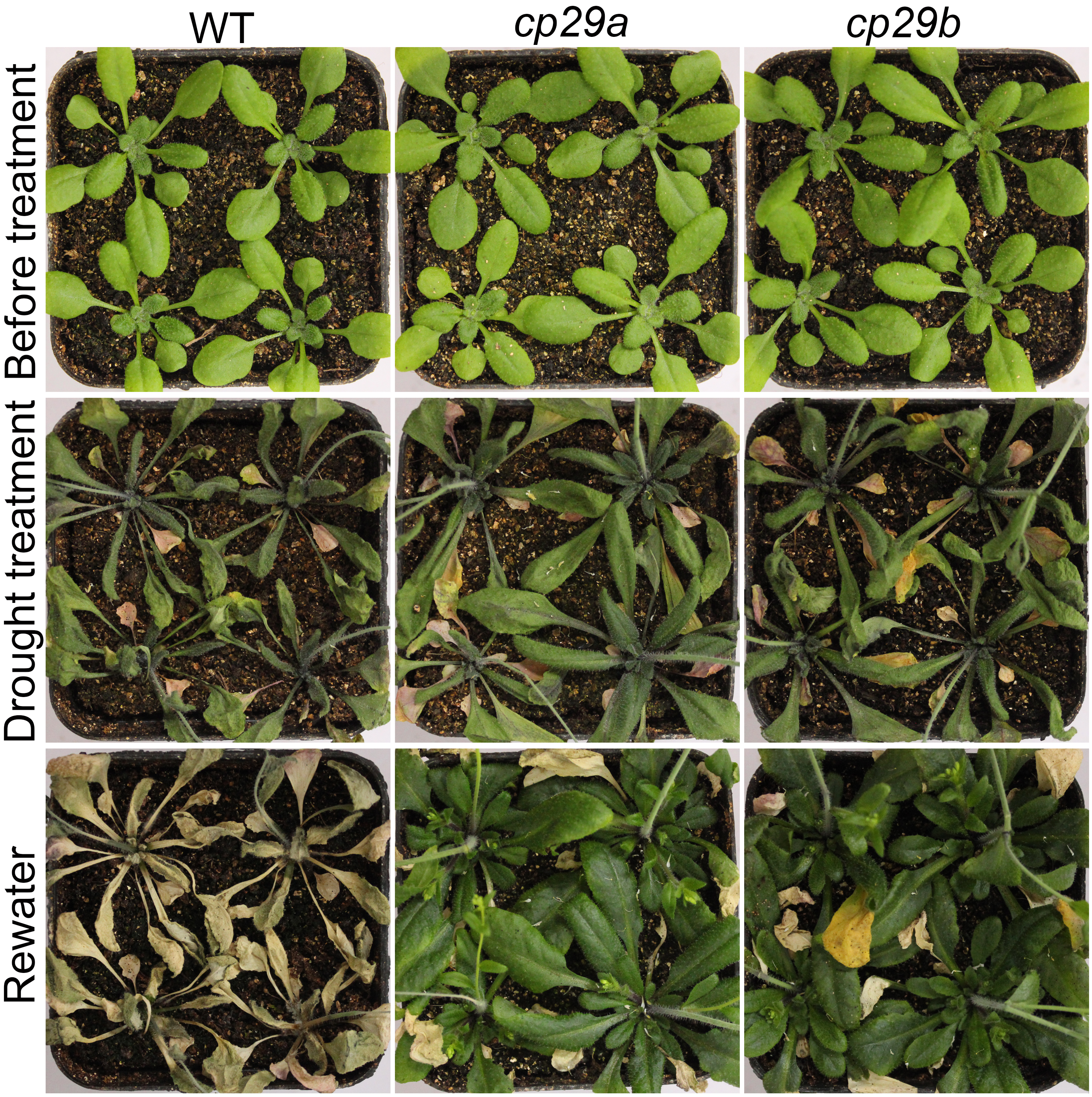

**Fig. S2** The impact of CP29A and CP29B on plant drought tolerance. Three-week-old well-watered WT, *cp29a*, and *cp29b* seedlings were subjected to drought stress by stopping watering for 16 days and then were rewatered to allow recovery for 5 days.

**Fig.S3** The impact of *DRIR*, CP29A, and CP29B on leaf bleaching under 4℃ conditions. Seven-day-old seedlings of WT, *drir^D^*, *DRIR* OE, *cp29a*, and *cp29b* grown under normal conditions were transferred to a 4°C growth chamber for an additional one-month growth period.

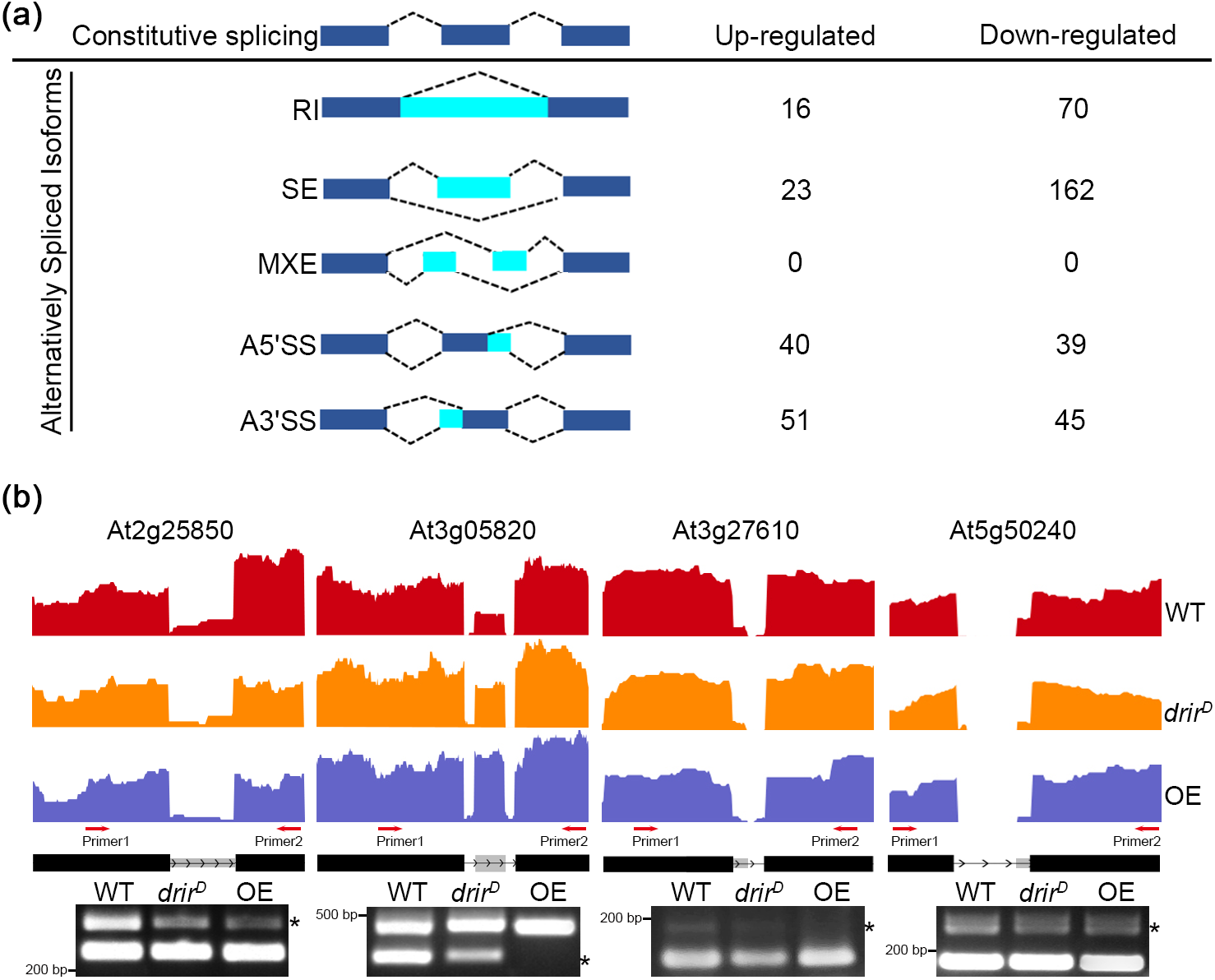

**Fig.S4** *DRIR* affects pre-mRNA alternative splicing under drought stress. (a) Types of different-type alternative splicing events shared by *drir^D^* and *DRIR* OE. Constitutive splicing refers to mRNA exon constitutive splicing. (b) The IGV gene tracks and RT-PCR validations of representative RI (At2g25850), SE (At3g05820), A5’SS (At3g27610), and A3’SS (At5g50240) events in WT, *drir^D^*, and *DRIR* OE seedlings. The IGV gene tracks display sequencing signal intensities for representative alternative splicing events in the transcriptomes. Exons are shown as black boxes, introns are represented as lines and the alternative regions are indicated by gray boxes. Red arrows indicate the positions of designed primers. Asterisks represent the alternative spliced isoforms.

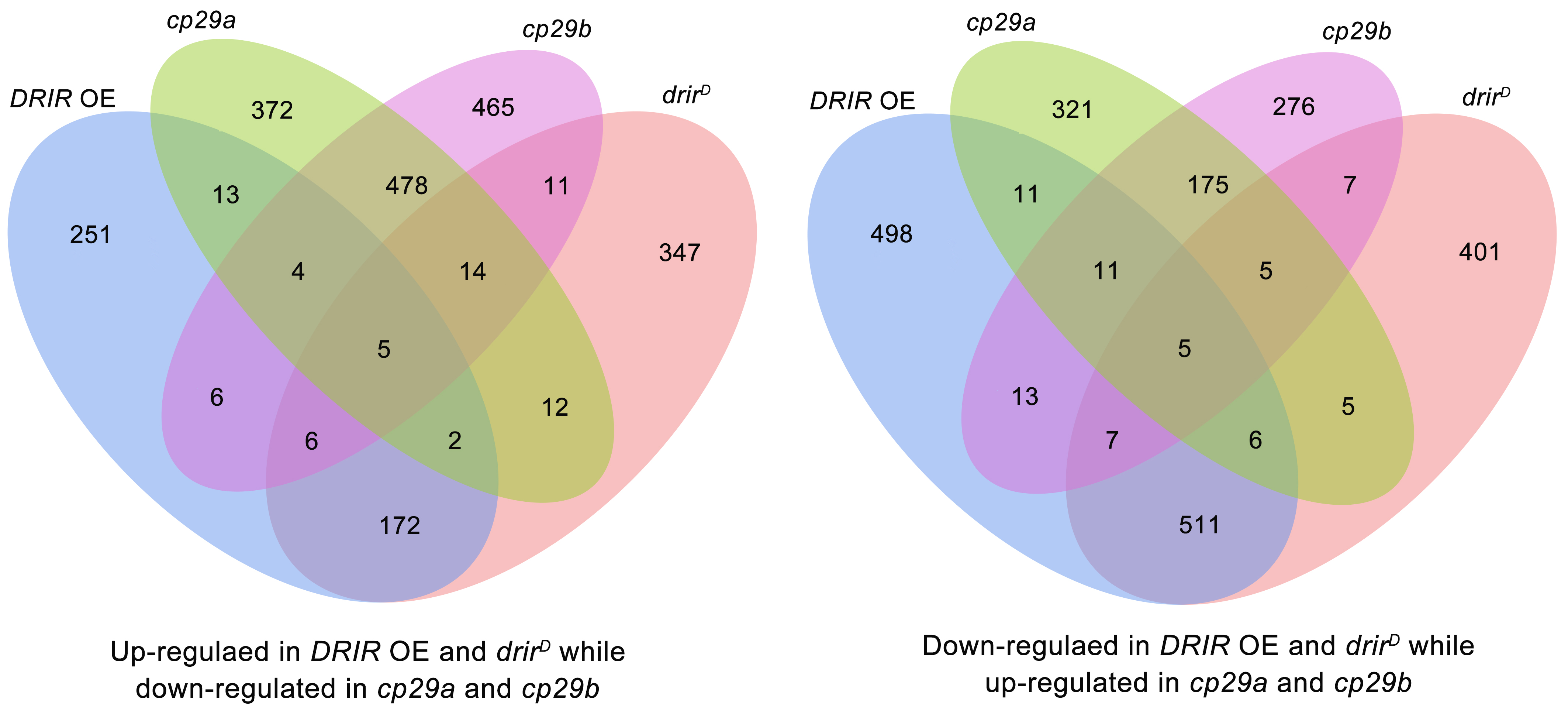

**Fig. S5** Venn diagrams showing the overlaps among differential splicing events that were up- or down-regulated in *drir^D^* and *DRIR* OE but were inversely regulated in *cp29a* and *cp29b*.

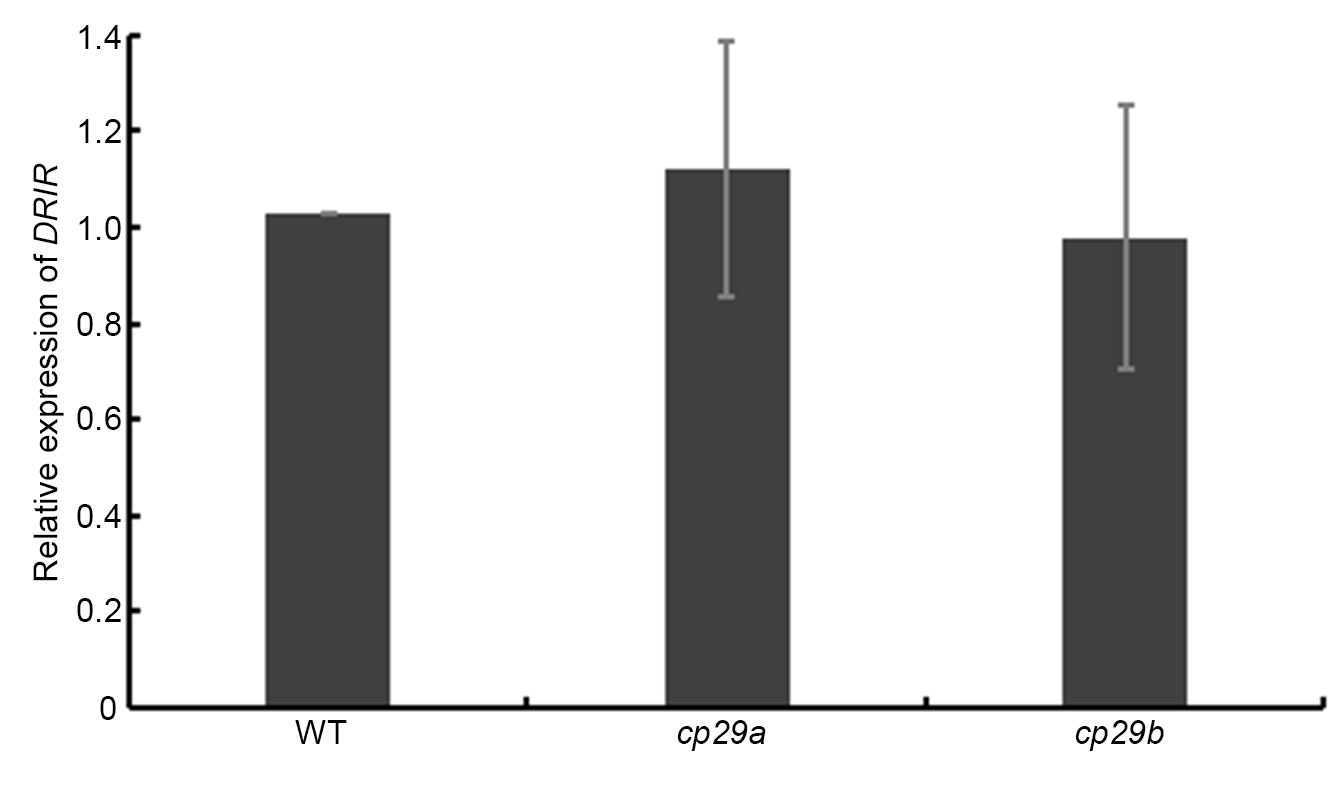

**Fig. S6** Transcript levels of *DRIR* inWT, *cp29a*, and *cp29b* seedlings. Data represent means ± SD from three biological replicates.

| **Table S1. Primers used in this study** | |
| --- | --- |
| **Primer name** | **Sequence** |
| DRIR-RNAi-F1 | TACATTTACAATTACCATGGGAGGACACGTCAGACGAAG |
| DRIR-RNAi-R1 | CGATTGGGCGCGCCCCAAACTCCAAACTCCTTTATTTC |
| DRIR-RNAi-F2 | GGATCCTAGGTGAGTCAAACTCCAAACTCCTTTATTTC |
| DRIR-RNAi-R2 | GTCTTAATTAACTCTGAGGACACGTCAGACGAAG |
| CP29A OE-F | TCTAGAATGTCTGCCTCTGCTTCTTC |
| CP29A OE-R | GGTACCAAATTGGCCTCTTGGTGGC |
| CP29B OE-F | TCTAGAATGGCTGCTTCAGCTTCG |
| CP29B OE-R | GGTACCATATTGGCGCCTTGGAGGC |
| DRIR SP6-F | ATTTAGGTGACACTATAGATGCAATTTGTGTAAACAATCTCGTGTG |
| DRIRSP6-R | CAAACTCCAAACTCCTTTATTTCTTAACCAAAAG |
| GST-CP29A-F | GGATCCATGTCTGCCTCTGCTTCTTC |
| GST-CP29A-R | GTCGACTCAAAATTGGCCTCTTGGTGGC |
| GST-CP29B-F | GGATCCATGGCTGCTTCAGCTTCG |
| GST-CP29B-R | GTCGACTCAATATTGGCGCCTTGGAGGC |
| GST-GRP7-F | GGATCCATGGCGTCCGGTGATGTTGAG |
| GST-GRP7-R | GTCGACTTACCATCCTCCACCACCAC |
| 18S rRNA-F | CGGCTACCACATCCAAGGAA |
| 18S rRNA-R | GCTGGAATTACCGCGCGGCT |
| DRIR-RT/RIP-qPCR-F | ATTTGCGGTGAGCGTGGATC |
| DRIR-RT/RIP-qPCR-R | TGTCCCCAAGCTCTATTGCAGG |
| AT1G23230 AS/RIP-F | gttcctagcattgcaatggttga |
| AT1G23230 AS/RIP-R | ggcttgcttagcagagaagca |
| AT1G23230 AS-R1 | agcatttcgctgggccaatt |
| AT4G08170 AS/RIP-F | caaattggaaaagaatggcgtcga |
| AT4G08170 AS/RIP-R | tccgcaacacactgaagcat |
| AT4G08170 AS-R1 | gactgacgattacggagatgtagt |
| AT1G62660 AS/RIP-F | ctattttgtcgtggcaacgcac |
| AT1G62660 AS-R1 | cttgttcctcaagttatggtctcat |
| AT1G62660 AS/RIP-R | tgtcaccccacacagctgca |
| AT5G04980 AS/RIP-F | catgccatcctttgaacactcga |
| AT5G04980 AS-R1 | cacaatcggttcctttcctccca |
| AT5G04980 AS/RIP-R | acgcttgggacactcttctca |
| AT2G17840 AS/RIP-F | gcgcatatcctgctgcatcga |
| AT2G17840 AS-R1 | tcgttgatcacaagtaagtaaaatcgagt |
| AT2G17840 AS/RIP-R | acaaccgaactcgttgatcaca |
| AT5G51820 AS/RIP-F | agtaacagcatgcatggcatca |
| AT5G51820 AS-R1 | cgactttgatctcatcagaggtct |
| AT5G51820 AS/RIP-R | ttccgactatttggagctcatgga |
| Actin-F | CGACAATGGAACTGGAATGGTTA |
| Actin-R | GTGCCTCGGTAAGTAGAATAGGA |
| CP29A protoplast -pEZL-NL-F | ACACGCTCGAGCTCAATGTCTGCCTCTGCTTCTTC |
| CP29A protoplast -pEZL-NL-R | GGGCCCGCGGTACCGTAAATTGGCCTCTTGGTGGC |
| CP29B protoplast -pEZL-NL-F | ACACGCTCGAGCTCAATGGCTGCTTCAGCTTCG |
| CP29B protoplast -pEZL-NL-R | GGGCCCGCGGTACCGTATATTGGCGCCTTGGAGGC |
